# Engineering microvascular networks using a KLF2 reporter to probe flow-dependent endothelial cell function

**DOI:** 10.1101/2023.10.31.565021

**Authors:** Adriana Blazeski, Marie A. Floryan, Oscar R. Fajardo-Ramírez, Elamaran Meibalan, Jesús Ortiz-Urbina, Emmanouil Angelidakis, Sarah E. Shelton, Roger D. Kamm, Guillermo García-Cardeña

## Abstract

Shear stress generated by the flow of blood in the vasculature is a potent regulator of endothelial cell phenotype and vascular structure. While vascular responses to flow are complex and context-dependent, endothelial cell signaling in response to shear stress induced by laminar flows is coordinated by the transcription factor KLF2. The expression of KLF2 in endothelial cells is associated with a quiescent, anti-inflammatory phenotype and has been well characterized in two-dimensional systems, but has not been studied in three-dimensional *in vitro* systems. Here we develop engineered microvascular networks (MVNs) with a KLF2-based endothelial cell sensor within a microfluidic chip, apply continuous flow using an attached microfluidic pump, and study the effects of this flow on vascular structure and function. We found that culture of MVNs exposed to flow for 48 hours that resulted in increased expression of the KLF2-GFP-reporter display larger vessel diameters and decreased vascular branching and resistance. Additionally, vessel diameters after the application of flow were independent of initial MVN morphologies. Finally, we found that MVNs exposed to flow have improved vascular barrier function and decreased platelet adhesion. The MVNs with KLF2-based flow sensors represent a powerful tool for evaluating the structural and functional effects of flow on engineered three-dimensional vascular systems.

## Introduction

Models of human microvascular physiology have been instrumental in bridging the gap between two-dimensional *in vitro* systems and animal models for studies of vascular biology and mechanisms of disease. In particular, self-assembled microvascular networks (MVNs) can recapitulate tissue-specific heterocellular interactions, complex three-dimensional geometries, and cell-matrix signaling^1^ to mimic the structure and function of *in vivo* vascular beds. These systems have been used to model a number of aspects of vascular biology, including vasculogenesis^2–4^, effects of interstitial flow on vascular formation^5^, and transport of biologics across the vascular barrier^6^. Additionally, MVNs with tissue-specific cells have been used to model biological environments like the blood-brain barrier^7^ and metastasizing tumors^8^.

A key element that can further improve engineered MVNs as physiological representations of human vasculature is the incorporation of flow. In vivo, vessels are constantly exposed to hemodynamic forces generated by the flow of blood, which regulates a host of processes during vascular development and homeostasis^9,10^. In particular, shear stress, the frictional force per unit area tangential to the endothelial surface, varies across vessels in the body from 0.1 Pa to 5 Pa^11^, and has been shown to regulate vascular remodeling, homeostasis, and cell fate determination^12^. Importantly, these processes are highly dependent on the direction and magnitude of shear stress in the vasculature^13^. Additionally, it has been proposed that endothelial cells (ECs) are tuned to a range of shear stresses that promote homeostasis, activate an inflammatory phenotype when exposed to shear stresses outside of this range, and will remodel vascular diameter in order to return to this setpoint^14^.

Because of this, readouts of vascular physiology in response to flow are needed to indicate the presence of threshold levels of shear stress and a corresponding effect on EC phenotype. Expression of Krüppel-like factor 2 (KLF2), a transcription factor that has been identified as a key integrator of endothelial response to shear stress^15–18^, is ideally suited as a cell-based readout of flow. The expression of KLF2 is induced in ECs by laminar flow^19^ and has been shown to regulate pathways involved vessel development, inflammation, thrombosis, and vascular tone^15^. Its expression is associated with a quiescent, vasoprotective endothelial phenotype that has been extensively characterized^15,17,20^. Importantly, a cellular KLF2 reporter system has been developed and shown to recapitulate key aspects of the expression of the endogenous gene^21^.

In this study, we incorporate a KLF2-based EC sensor into MVNs within a microfluidic system and apply circulating, continuous, unidirectional flow. We show that shear stresses activate the KLF2 reporter in ECs within a complex, three-dimensional network of vessels and respond by altering vascular geometry, including increasing vascular diameter. Additionally, flow conditioning improves the barrier function of MVNs and reduces platelet adhesion in the vessels. We further show that vessels with different starting geometry will remodel to a similar geometric endpoint in response to flow. Our system represents an engineering approach to microvascular models with flow that incorporates a built-in cellular flow reporter system. Importantly, it provides a useful readout that shear stress magnitude and timing has reached levels sufficient to elicit the KLF2-associated phenotype in MVNs and establishes a baseline for application of flow in other engineered systems.

## Results

### Microfluidic system with engineered vasculature containing flow-responsive endothelial cell reporter and *on chip* pump

Our microfluidic platform includes a chip with a central channel that houses the engineered MVNs bordered by media channels that are connected directly to a pump^22^ (**Fig. 1a**). This *on chip* pump is composed of two layers of PDMS with a silicone membrane between them that can be actuated at the pumping chamber to displace fluid and move cell culture media through the system. Two fluid capacitors are used to establish a pressure difference that drives flow across the microfluidic device while two check-valves are incorporated to ensure unidirectional flow. The pump is operated by applying cyclical, positive air pressure at the pumping chamber under the control of a solenoid. The pump inlet and outlet ports interface with the media channels bordering the MVNs. MVNs are formed by encapsulating human umbilical vein endothelial cells (HUVECs) and human lung fibroblasts into a fibrin gel and allowing them to self-assemble into lumenized vessels over the course of 4 to 5 days^23,24^. ECs incorporated a reporter based on GFP expression driven by the human KLF2 promoter^21^, allowing them to act as flow sensors in the system (**Fig. 1a**). At 5 days of culture, MVNs are fully perfusable with dextran (**Fig. 1b**), indicating that they have formed a connected bed of patent vessels through which media can be circulated. EC containing the KLF2 reporter and cultured in two-dimensional monolayers responded to application of laminar flow by turning on GFP fluorescence (**Supplementary Fig. 1**) and we subsequently tested whether these cells could act as flow sensors in the three-dimensional MVNs. Application of flow using the PDMS pump for 18 hours resulted in activation of GFP localized to vessel walls in MVNs cultured under flow (“flow MVNs”) but not in MVNs cultured under static conditions (“static MVNs”) (**Fig. 1c**). Static and flow MVNs were subsequently dissociated into single cells and stained for VE-cadherin before being separated into EC (VE-cadherin^+^) and fibroblast (VE-cadherin^-^) populations using flow cytometry (**Fig. 1d, Supplementary Fig. 2**). While fibroblasts dissociated from both static and flow MVNs had similarly low expressions of fluorescence, EC from flow MVNs exhibited an increase in GFP expression compared to those from static MVNs (**Fig. 1e**). Quantification of GFP expression in sorted ECs indicated an increase in GFP-positive cells from 12.0% in static MVNs to 56.2% in flow MVNs (**Fig. 1f**). Additionally, ECs from flow MVNs exhibited increased KLF2 gene expression compared to ECs from static MVNs (**Fig. 1g**, n=5 pooled devices). Altogether, these data indicate that our system is suitable for applying flow to 3-dimensional MVNs and that KLF2-GFP serves as an EC-specific flow reporter that is associated with the induction of key flow-induced gene transcripts.

**Figure 1.**
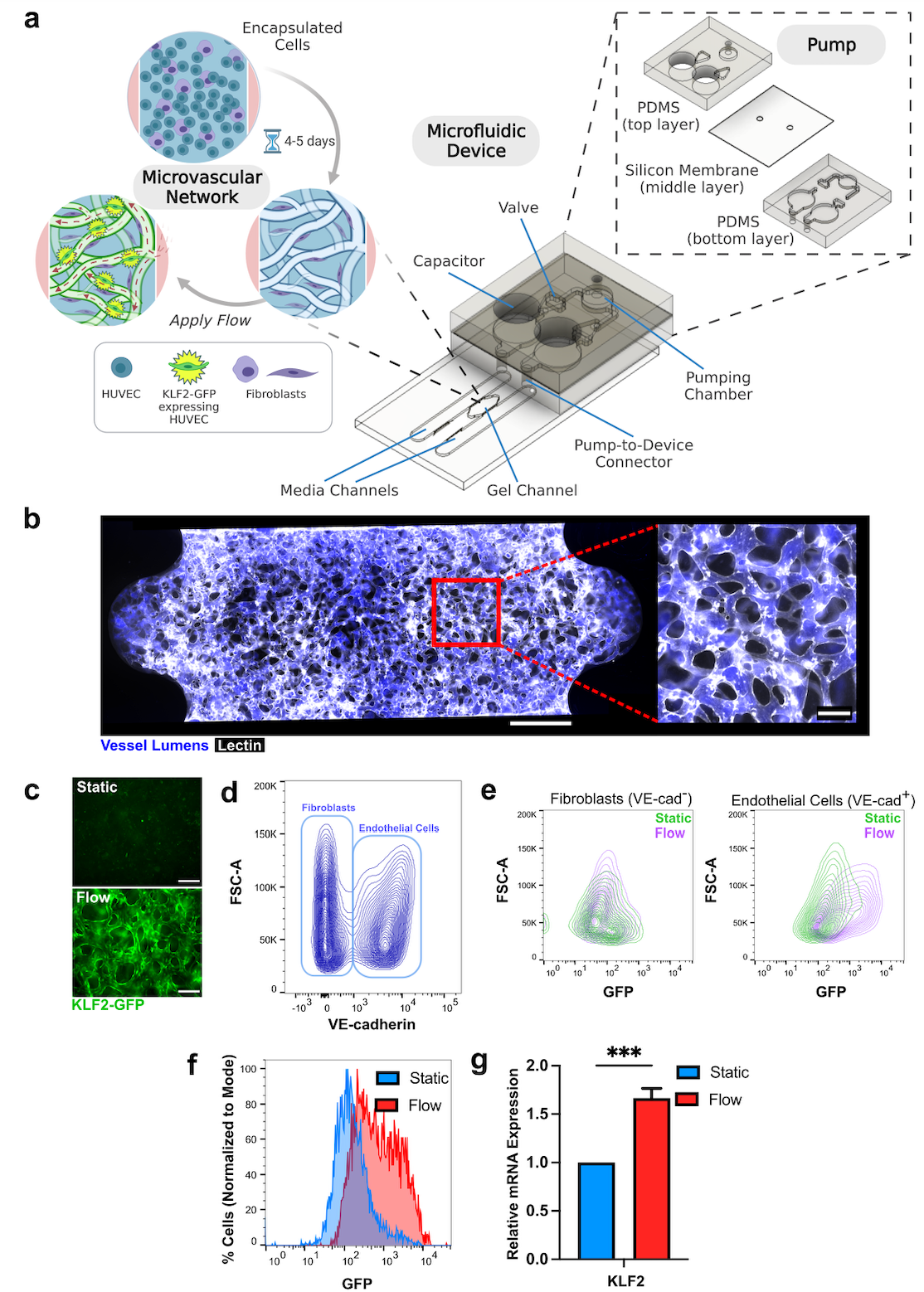
Activation of KLF2-GFP reporter in MVNs with flow. **a** Schematic of microfluidic device and attached pump. HUVECs expressing KLF2-GFP and fibroblasts are seeded into the gel channel and self-assemble into MVNs. **b** Example of MVN stained with lectin (white) and perfused with fluorescent dextran (blue). Scale bar = 500 μm in full device image. Scale bar = 100 μm in close-up image of vasculature. **c** Image of KLF-GFP (green) expression in MVN after application of flow for 18 hours in MVN after culture in static conditions. Scale bars = 200 μm. **d** Flow cytometry analysis of cells in MVNs labelled with VE-cadherin. VE-cadherin^+^ endothelial cells and VE-cadherin^-^ fibroblasts are indicated by bounding rectangles. **e** Flow cytometry analysis of each population from MVNs cultured under static (green) and flow (purple) conditions. **f** Percentage of endothelial cells expressing GFP under static and flow conditions. **g** KLF2 mRNA expression in endothelial cells extracted from MVNs and separated out using flow cytometry as in **d** (measurement from 5 pooled MVNs). *** indicates p<0.001.

**Figure 2.**
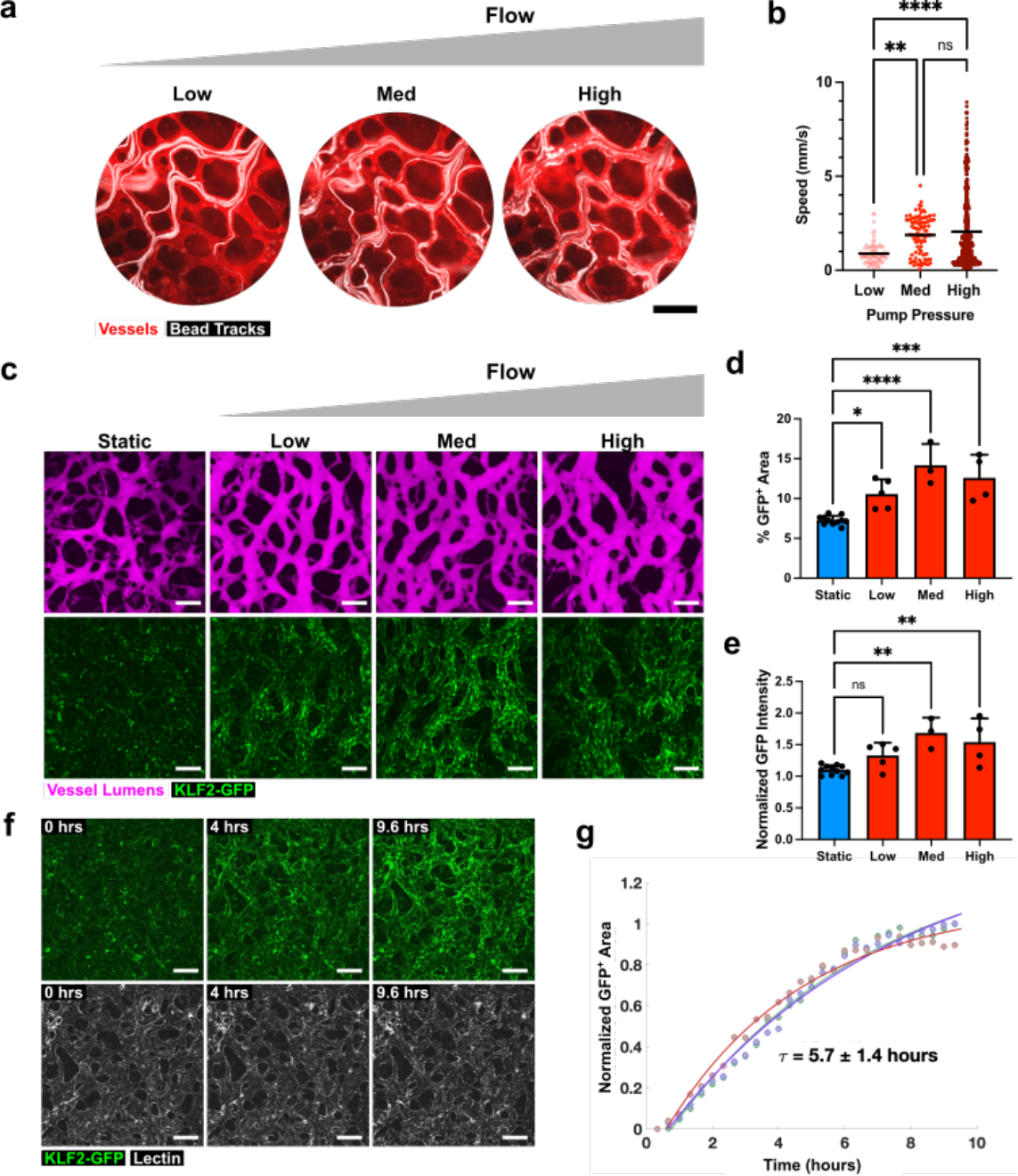
Characterization of flow levels and time course in engineered MVN system. **a** Paths of fluorescent beads (white) in MVN (stained with red lectin) as pressure is increased in the pumping chamber. Scale bars = 200 μm. Each image represents a projection of bead location over 30 seconds. “Low”, “Med”, and “High”, correspond to application of 3 kPa, 6 kPa, and 9 kPa pressure, respectively, in the pumping chamber. **b** Average bead velocities for range of pressures in the pumping chamber, calculated from images of bead velocities (n = 3 samples, with 14-45 bead paths per sample). **c** Examples of MVNs, visualized after perfusion with fluorescent dextran (magenta), and their KLF2-GFP expression (green) under static conditions or after 48 hours of flow at the range of flow rates characterized in b. Scale bars = 200 μm. **d** Percent of total imaging area with KLF2-GFP signal. **e** KLF2-GFP signal intensity normalized to lowest intensity static sample. **f** Time series of z-stack confocal image projections of MVNs after the start of flow corresponding to medium pressure and acquired every 20 minutes. Samples were stained with lectin (white) to outline vessels. Scale bars = 200 μm. **g** Time course of KLF-GFP expression in MVNs after initiation of flow. GFP^+^ area for different regions in the MVN was modeled by an exponential fit and an average time constant for KLF2-GFP activation was calculated. τ is reported as mean ± standard deviation for 3 regions of interest. Data in **b, d**, and **e** reports mean and standard deviation from n = 3-11 MVNs. * indicates p<0.05, ** indicates p<0.01, *** indicates p<0.001, **** indicates p<0.0001, and ns indicates p>0.05.

### Characterization of flow levels and timing in engineered system

Having determined that the KLF2-GFP reporter system can act as a flow sensor in MVNs, we next sought to determine the levels of flow needed to activate the GFP signal. To this end, we introduced fluorescent beads into our MVN and pump system and captured videos of their trajectories in the vessels in response to increasing pumping flow rates, achieved by increasing maximum air pressures applied at the pumping chambers. Bead positions were superimposed over time to generate trajectories (**Fig. 2a**), which were subsequently used to calculate average bead velocities within MVNs (**Fig. 2b**). Pressures of 3 kPa (“Low”), 6 kPa (“Medium”), and 9 kPa (“High”) applied at the pumping chamber resulted in bead velocities of 0.8-1.02 mm/s, 1.7-2.1 mm/s, and 1.9-2.2 mm/s, 95% confidence intervals of the means respectively, (**Fig. 2b**, n = 14-45 trajectories in n = 3 MVNs). The three levels of flow applied in our system all elicited KLF2-GFP expression in the MVNs (**Fig. 2c**). The percentage of imaging area positive for KLF2-GFP signal was significantly increased in all flow groups relative to static controls (7.3±0.6%, 10.6±1.8%, 14.2±2.7%, and 12.6±2.9% for static, low, medium, and high flow, respectively, where n = 3-11 per group), but did not differ significantly between flow groups (**Fig. 2d**). Normalized GFP intensity, indicative of the level of KLF2 expression in MVNs, was elevated significantly only in the medium and high flow groups compared to static controls (measuring 1.1±0.1 for static, 1.3±0.2 for low, 1.7±0.2 for medium and 1.5±0.4 high flow, respectively, where n = 3-11 per group), but did not differ significantly between these two flow groups (**Fig. 2e**). Based on these observations, we determined that 6 kPa of pressure applied at the pumping chamber, corresponding to “medium” flow, was sufficient to activate KLF2-GFP in our system and we used this level of flow for the remainder of the experiments in our study.

We next characterized the time course of KLF2-GFP activation in MVNs under flow by acquiring fluorescent confocal images across the volume of vessels over the course of nearly 10 hours of flow (**Fig. 2f, Supplementary Video 1**). Calculating the GFP-positive area from two-dimensional projections of these images and applying an exponential fit to the time course yielded a time constant for GFP expression of 5.7±1.4 hours (**Fig. 2g**, n=3). The 48 hours of flow applied in all subsequent experiments, therefore, were sufficient to ensure robust KLF2-GFP expression.

We used a computational approach^25^ to determine the magnitudes of the wall shear stresses within the MVNs exposed to flow (**Fig. 3**). To accomplish this, we constructed a network of vascular branches based on confocal images spanning the width of individual MVNs and applied a pressure difference across this network. For each MVN, we computed the pressure difference across the network that would result in an average flow speed of 2 mm/s in the MVN (**Fig. 3d**), mimicking our bead measurements under medium flow (**Fig. 2b**) and simulating the pressure difference achieved by the pump. This revealed preferential flow paths in the MVNs that were characterized by higher flow rates (**Fig. 3a**). The flow speeds (**Fig. 3b**) and wall shear stresses (**Fig. 3c**) varied across the branches within each MVN, but the average shear stress within a MVN was 0.53±0.05 Pa (**Fig. 3e**, n=5).

**Figure 3.**
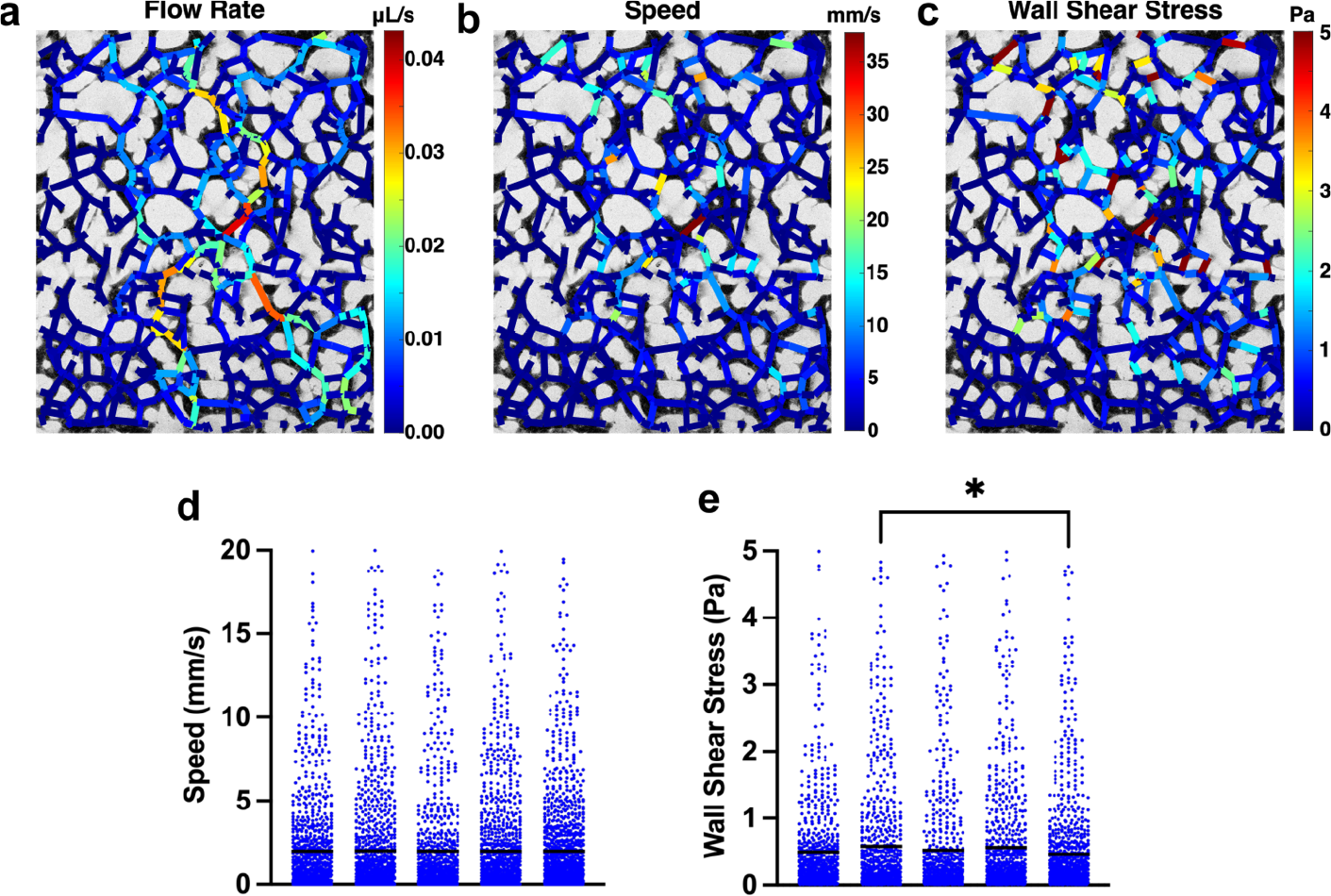
Estimates of wall shear stress in MVNs. **a** Map of flow rate in each branch of MVN. **b** Map of speed in each branch of MVN. **c** Map of wall shear stress in each branch of MVN. **d** Flow speeds in each branch of 5 individual MVNs. **e** Wall shear stresses in each branch of 5 individual MVNs. Each point in **d** and **e** represent an independent vessel branch and each column represents an independent MVN. Horizontal black lines indicate mean value for the MVN. * indicates p<0.05. All other comparisons between MVNs in **d** and **e** were not statistically significant.

### Changes in vascular morphology in response to flow

Because our system incorporates three-dimensional MVNs, it provided us with an opportunity to determine the effects of flow on vascular geometry. For these studies, we applied flow to MVNs for 48 hours and compared them to control MVNs cultured under static conditions. Fluorescent dextran perfused into MVNs was used to delineate the vascular lumens and to segment out vessels for morphologic analyses (**Fig. 4a**). Flow MVNs were characterized by strong KLF2-GFP expression along vascular walls (**Fig. 4a**) with corresponding increases in percent area positive for GFP (6.3%±2.4%, n=8 static versus 12.3%±3.4%, n=6 flow, **Fig. 4b**) and in normalized GFP intensity (1.1±0.1, n=8 static versus 1.7±0.3, n=7 flow, **Fig. 4c**) when compared to static MVNs. While the total area taken up by vessels, calculated from maximum intensity projections of images over the volume of dextran-perfused vessels, did not differ between static and flow MVNs (1.0±0.1 mm^2^, n=12 for static and 1.1±0.1 mm^2^, n=11 for flow, **Fig. 4d**), other aspects of branch geometry differed significantly between the two groups. Flow MVNs had a significantly lower number of branches (108±14, n=12 for static and 87±12, n=11 for flow, **Fig. 4e**), and a significantly greater average branch length (67.7±8.1 μm, n=12 for static and 77.3±7.6 μm, n=11 for flow, **Fig. 4f**) compared to static MVNs. Interestingly, cross-sectional views of flow MVNs revealed larger vessel lumens than those in static MVNs (**Fig. 4g**). Flow was associated with a significant increase in average vessel diameter (67.1±12.5 μm, n=12 for static and 96.8±21.1μm, n=11 for flow, **Fig. 4h**) and a wider distribution of vessel diameters (**Fig. 4i**).

**Figure 4.**
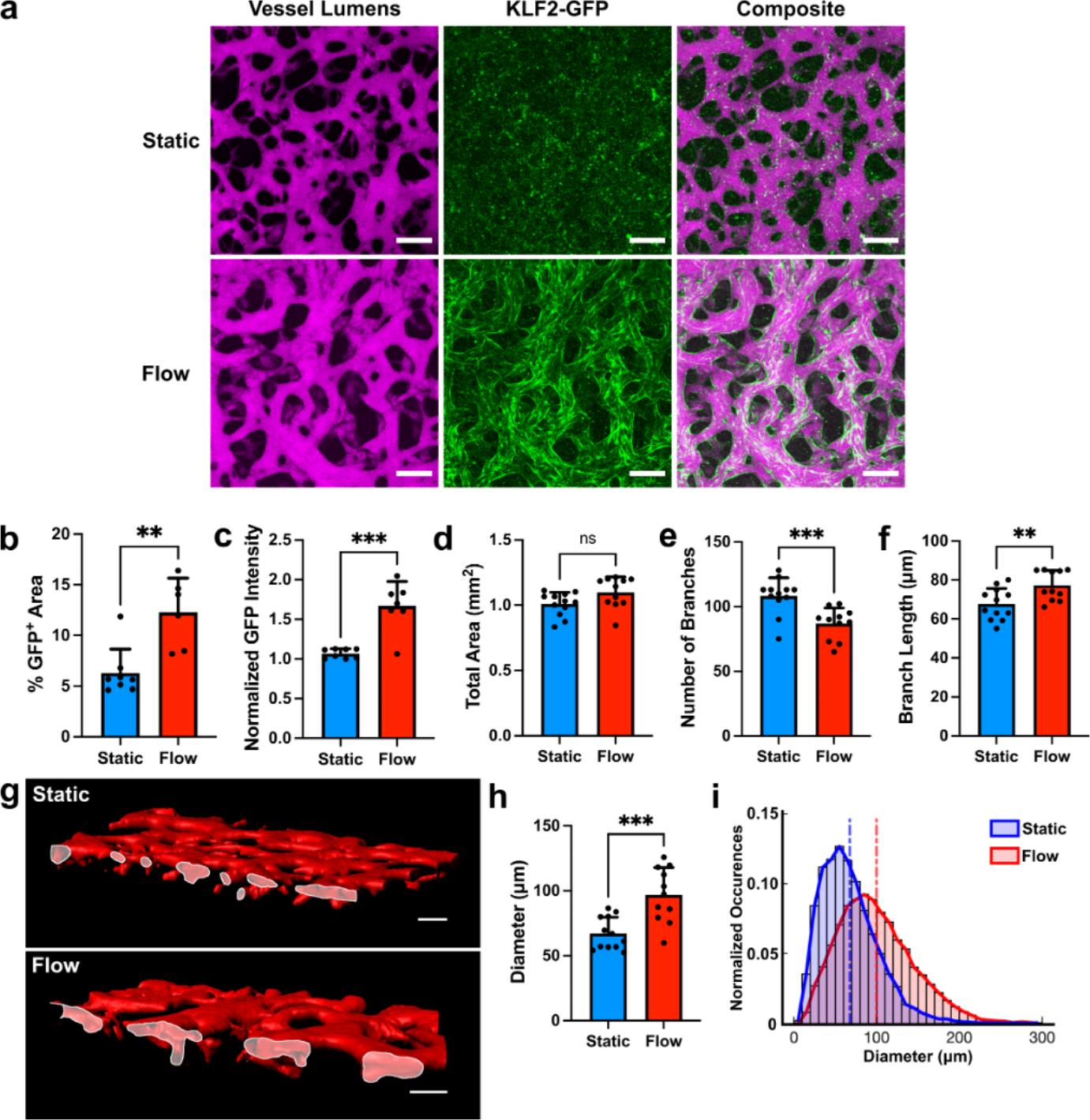
Flow-induced changes in vascular geometry. **a** MVNs cultured under static (top) or flow (bottom) conditions. Vessels were perfused with fluorescent dextran (magenta) to visualize the intravascular space. MVNs cultured under flow expressed the KLF2-GFP reporter (green). Scale bars = 200 μm. **b** Percent of total imaging area with KLF2-GFP signal. **c** KLF2-GFP signal intensity normalized to lowest intensity static sample. **d-f** Number of branches, total area covered by vessels, and average branch length in MVNs. **g** Surface reconstruction of static MVNs (top) and MVNs under flow (bottom) cut across the xz plane to show vessel openings (outlined in white). **h** Average branch diameter in MVNs. **i** Distribution of vascular diameters in MVNs cultured under static (blue) and flow (red) conditions. Vertical dashed lines indicate average diameters across all samples in each group. In panels **b-f** and **h**, n=11-12 MVNs per group. ** indicates p<0.01, *** indicates p<0.001, and ns indicates p>0.05.

### Effect of vascular geometry on flow response

We next sought to determine whether the morphological remodeling of vessels under flow was affected by the initial geometry of the MVNs. The average vessel diameter in MVNs was varied by changing the concentration of thrombin that was mixed with fibrinogen prior to gel formation: 1.8 U/mL thrombin and 1.4 U/mL thrombin were used to make smaller and larger vessels, respectively. Decreasing thrombin concentration increased the gelling time, allowing the cells more time to settle in a smaller volume, and resulting in large vessels (**Fig. 5a**). This resulted in the generation of MVNs with an average diameter of 52.3±1.7 μm (smaller vessels) or 59.8±3.2 μm (larger vessels, **Fig. 5e**). Application of flow in both larger and smaller vessels increased the percent area where KLF2-GFP was expressed (5.6±0.8 %, n=4 for smaller static vessels, 16.5±3.2 %, n=4 for smaller flow vessels, 6.7±0.7 %, n=4 for larger static vessels, and 14.7±2.8 %, n=8 for larger flow vessels), and the percent GFP-positive area was not significantly different in either flow group (**Fig. 5b**). However, normalized GFP intensity was only significantly increased in response to flow in the MVNs with smaller vessels (1.1±0.04, n=4 for smaller static vessels, 1.5±0.1, n=4 for smaller flow vessels) and did not change significantly in response to flow in MVNs with larger vessels (1.0±0.1, n=4 for larger static vessels, 1.2±0.2, n=8 for larger flow vessels, **Fig. 5c**). Permeability, a measure of vascular barrier function, tended to decrease with application of flow in both smaller and larger vessels, but this effect was not statistically significant (**Fig. 5d**), although larger flow vessels had significantly lower permeability than static smaller vessels (6.6x10^-8^±4.0x10^-8^ cm/s, n=4 for smaller static vessels, 1.6x10^-8^±7.0x10^-9^ cm/s, n=8 for larger flow vessels). Finally, vessel diameters increased in response to flow in both smaller and larger vessels (52.3±1.7 μm, n=4 for smaller static vessels, 67.5±2.4 μm, n=4 for smaller flow vessels, 59.8±3.2 μm, n=4 for larger static vessels, and 70.4±3.4 μm, n=4 for larger flow vessels, **Fig. 5e**). Interestingly, diameters of smaller and larger vessels under flow were not significantly different, indicating that flow can be used to achieve a similar vessel diameter endpoint, regardless of the starting vessel diameter.

**Figure 5.**
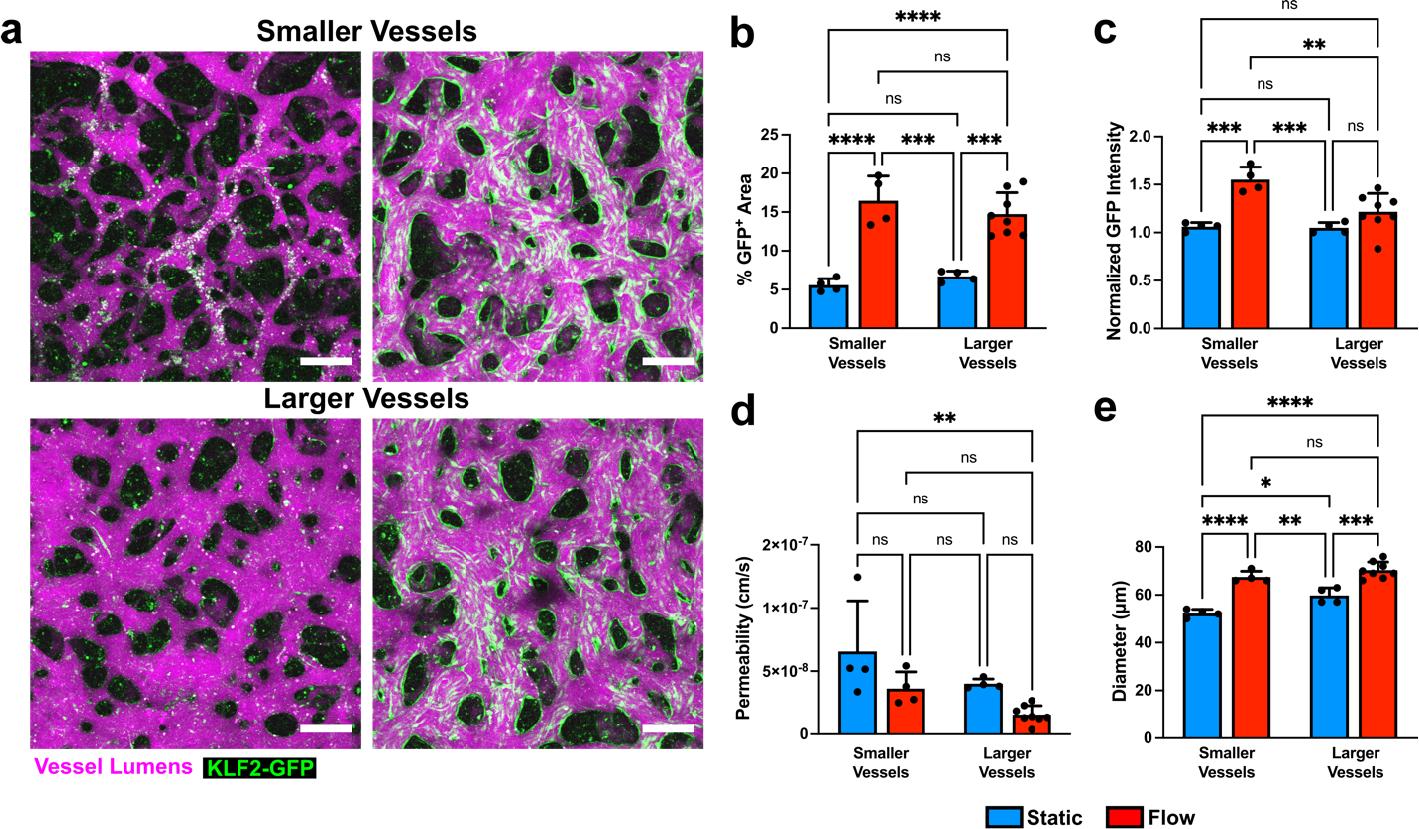
Modulation of vascular responses to flow by initial vascular geometry. **a** MVNs with smaller (top) or larger (bottom) average diameters perfused with fluorescent dextran (magenta) under static culture and after culture under flow that activates KLF2-GFP expression (green). Scale bars = 200 μm. **b-e** Percent of total imaging area with KLF2-GFP signal, KLF2-GFP signal intensity normalized to lowest intensity static sample, vascular permeability, and average vascular diameter in smaller and larger vessels under static culture (blue) and after culture under flow (red). n=4-8 MVNs in each group of panels **b-e**. * indicates p<0.05, ** indicates p<0.01, and **** indicates p<0.0001.

### Changes in vascular function in response to flow

Alongside of changes in vascular geometry, we characterized the effects of flow on several functional properties of the MVNs. To measure vascular barrier function, we perfused MVNs with fluorescent dextran, acquired images of the networks and their surrounding gel at 0 and 12 minutes, and measured the change in extravascular fluorescence over time (**Fig. 6a**). This rate of change in fluorescence, indicative of the rate of transport of dextran from the intra- to the extra-vascular space, was used to calculate vascular permeability^6^, which was significantly decreased in flow MVNs (9.6x10^-8^ ±6.4x10^-8^ cm/s, n=12 for static and 4.9x10^-8^ ±2.0x10^-8^ cm/s, n=11 for flow, **Fig. 6b**). In addition to improving vascular barrier function, the application of flow in MVNs significantly decreased hydraulic vascular resistance across the entire MVN (5.2 x10^13^ ±1.6 x10^13^ Pa s/m^3^, n=6 for static and 3.8 x10^12^ ±2.2 x10^12^ Pa s/m^3^, n=5 for flow, **Fig. 6c**). Finally, we investigated the effects of flow on modulating the thrombogenicity of the vascular wall by labeling platelets with an antibody against CD41 in suspension and perfusing them into MVNs cultured under static conditions or into MVNs cultured under flow immediately after ceasing flow and removing the devices from the pump (**Fig. 6d**). Fluorescent confocal imaging revealed many more CD41^+^ platelets in static MVNs than in flow MVNs (**Fig. 6e**) and the percentage area covered by CD41^+^ platelets was significantly higher in static MVNs (5.7±1.8 %, n=6 for static and 1.6±0.3 %, n=5 for flow, **Fig. 6f**). Thus, culture under flow improved vascular barrier function, decreased vascular network resistance, and decreased thrombogenicity in MVNs.

**Figure 6.**
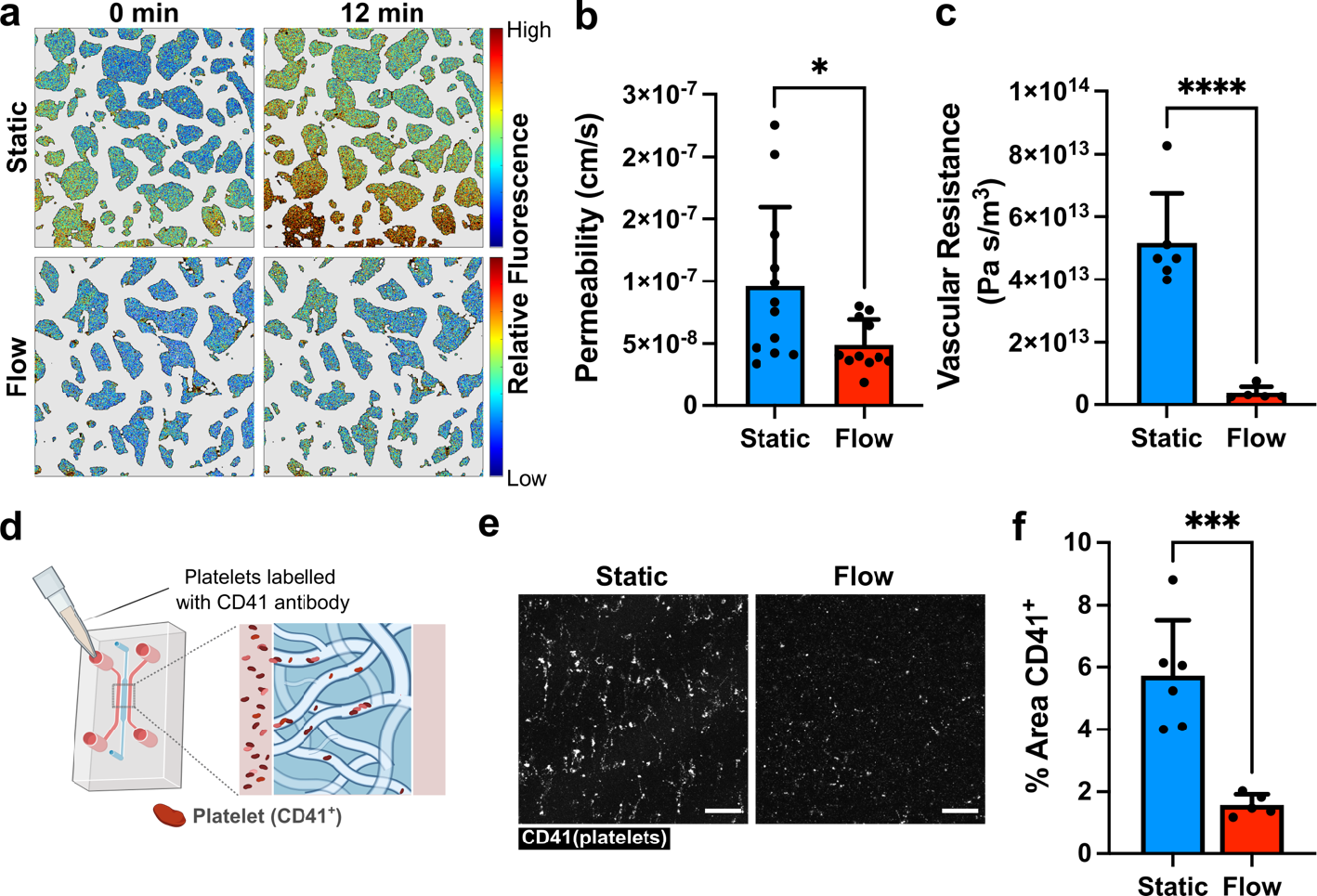
Flow-induced changes in vascular function. **a** Heat maps of extravascular fluorescence in MVN cultured under static (top row) and flow (bottom row) conditions immediately after vascular perfusion with fluorescent dextran (left) and after 12 minutes (right). Scale bars = 200 μm. **b** Vascular permeability in MVNs cultured under static conditions or after 48 hours of flow (n=11-12 per group). **c** Total vascular resistance of MVNs cultured under static conditions or after 48 hours of flow (n=5-6 per group). **d** Schematic of experimental design for introducing platelets into MVNs. **e** Volume projections of CD41-labelled platelets (white) in MVNs. Networks were cultured under static conditions or exposed to flow for 68 hours prior to addition of platelets in the networks. Scale bars = 200 μm. **f** Percent of image area covered by CD41-labelled platelets in MVN devices (n=5-6 per group). * indicates p<0.05, *** indicates p<0.001, and **** indicates p<0.0001.

## Discussion

The regulation of EC structure and function by hemodynamic forces^26,27^ makes the incorporation of flow an essential component in engineering physiologically-relevant models of the vasculature. In this study, we applied continuous, circulating flow to self-assembling MVNs in a microfluidic chip and studied the resultant changes in vascular geometry and function. Although many groups report systems with engineered vessels under flow, a majority of these constitute a single channel with a diameter over 100 μm^28–30^, and sometimes with a diameter of 500 μm or more^31,32^. Our self-assembled MVNs contain vessels with an average diameter of around 50 μm, resembling the size of smaller arterioles and venules in the body^33^. In systems similar to ours, which recapitulate a complex network of smaller-diameter vessels, flow has been achieved by applying hydrostatic pressure differences across the vascular networks^34,35^, which results in decreasing fluid flow as the pressures equilibrate over time. In contrast, our device drives continuous flow through the MVNs for sustained periods of time, which is necessary for the induction of flow-related genes in the ECs.

A key feature of our system is the incorporation of a flow-sensitive EC-specific reporter, driven by the promoter of KLF2^21^. While many genes are regulated by shear stress^9,36^, the choice of KLF2 expression as the basis for a flow sensor is particularly attractive because, within the vasculature, the expression of this transcription factor occurs only in ECs and is induced by sustained laminar flow^19,20^. In HUVECs, KLF2 induction is specifically tied to shear stress and is not induced by the application of cyclic strain^16^, allowing us to use it as a marker of a particular type of mechanical force. Importantly, KLF2 is a key orchestrator of endothelial gene expression programs that regulate blood vessel development, response to inflammation, thrombosis, and vascular tone^15^, and therefore serves as a signature of a well-defined endothelial phenotype. Given that differences in approach to applying flow in engineered systems can make it difficult to interpret findings across studies, reporters like the KLF2-GFP construct we use here can help verify whether the flow regimes applied produce a specific EC phenotype and are, therefore, comparable.

Additionally, we identified flow parameters sufficient for expression of KLF2 in our MVNs. In agreement with previous studies^37,38^, we found that two-dimensional EC monolayer preparations express KLF2 in a majority of cells in response to 0.5-1 Pa of shear stress. While the KLF2 reporter is activated in response to shear stresses starting at 0.2 Pa in monolayer cultures^39^, it had not previously been used in three dimensional vascular preparations. We found that applying flow for a minimum of 6 hours results in robust expression of KLF2 in MVNs, which can serve as a benchmark for other microphysiological vascular systems. The flows in our system, visualized by tracking the positions of fluorescent beads within the MVNs, appeared regular and laminar, consistent with unidirectional flows having little flow reversal that are known to activate KLF2^40^. Computational modeling of flow in our MVNs prior to flow-induced remodeling indicated an average wall shear stress of about 0.5 Pa, in the range of values reported for post-capillary venules^41^. At the same time, a range of wall shear stresses was present within each MVN, reflecting the heterogeneity of branch geometry and size within each device.

MVNs exposed to KLF2-inducing flows exhibited a marked increase in diameter. It has long been known that blood flow can regulate vascular diameter^42^ and outward remodeling has been associated with high shear stresses^43^. In the developing mouse, hemodynamic forces from blood flow are necessary for vascular remodeling^44^ and cause increases in vascular diameter by inducing vessel fusion in regions of high flow and EC migration towards regions with greater flow, rather than EC proliferation^45^. This type of vascular remodeling occurs when ECs experience shear stresses outside of a physiological set point^14^ and results in changes in diameter that ultimately restore shear stress levels to optimal, vasculature-stabilizing levels^27^.

We found that flow conditioning did not increase the range of flow diameters in the MVN system, but caused a rightward shift in the distribution of vessel diameters and resulted in longer and less-branched vessels throughout the MVNs. The presence of fewer but larger vessels along with the establishment of preferential flow paths as media was circulated through the MVNs, evident in our computational simulations of flow, suggests the occurrence of vascular pruning. This is a likely explanation for the widespread expression of KLF2-GFP in virtually all vessels after 48 hours of flow, since vessels along preferential paths were exposed to flow and activated the flow-responsive gene while other vessels were subjected to infrequent or low flow and regressed. Remodeling of vascular geometry in response to flow has been described *in vivo*, in multiple organisms. Chen et al. described hemodynamic regulation of vascular pruning during development of the zebrafish midbrain, where pruning was initiated in vascular segments with low and variable blood flow^46^. ECs from these pruning segments migrated to adjacent unpruned segments with higher flow, resulting in overall simplification of complex vascular networks over time^46^. Observations in the developing mouse retina have also revealed shear stress-mediated cellular polarization that directs migration toward high flow branches, resulting in destabilization and ultimate regression of low-flow branches and net movement toward and stabilization of high-flow branches^47^.

Cellular elongation and alignment along the direction of flow is often observed in planar EC preparations exposed to laminar shear stresses^48,49^. In our MVNs, ECs were highly elongated and oriented along the long axis of each vascular branch, even in static culture. This high degree of elongation and orientation was maintained with flow. Having established vascular remodeling and EC enlargement in our system over the course of 48 hours of flow, future work can focus on observing individual ECs within MVNs over time to determine the mechanisms of vessel widening and pruning.

In addition to structural changes, we observed numerous functional changes in MVNs cultured under flow. MVNs under flow had a significantly lower vascular resistance than those cultured under static conditions, in keeping with their overall larger vessel diameter. We also assessed vascular barrier function by perfusing fluorescent dextran through the MVNs and measuring changes in fluorescent intensity in the vascular and extravascular compartments ^6^ over time. MVNs under flow had a significantly decreased permeability, indicating a tighter endothelial barrier. This improvement in barrier function in response to flow is in line with previous studies. In straight channel vessels formed within collagen gels, flow improved vascular barrier function and extended the lifespan of vessels. In this system, improvements in barrier function and more organized and uniform expression of VE-cadherin at EC junctions were attributed to increased shear stresses and were not altered by changes in transmural pressure^50^. Further, it has been suggested that improvements in barrier function are due to shear stress-mediated activation of Rac and downstream actin filament reorganization^51^. In our studies, application of flow to improve the barrier function increases the physiological relevance of engineered MVNs. While it has been shown that three-dimensional MVN preparations have lower permeabilities than monolayer cultures, the further improvement in barrier function through the application of flow makes them similar to *in vivo* models for barrier studies^6^.

Haase and colleagues also applied circulating flow for 48 hours to complex MVNs^35^. While they report that continuous flow staves off changes in vessel geometries, like decreases in vascular diameter, observed in static cultures over time and causes a slight but non-significant improvement in vascular barrier function, our findings are more prominent: application of circulating flow in our system increases vascular diameter and significantly improves barrier function.

Finally, we assessed the anti-thrombogenic effect of KLF2 reporter-inducing flow in our MVNs. Our system was able to recapitulate this important aspect of flow, as MVNs cultured under flow exhibited adhesion of significantly fewer platelets than MVNs cultured under static conditions. It has been previously demonstrated that preconditioning EC monolayers with KLF2-activating flow decreased IL1-beta-dependent adhesion of leukocytes^15^ and our three-dimensional MVNs recapitulate this important anti-thrombogenic feature of flow-conditioning.

The KLF2-GFP EC sensor deployed in our system is a particularly useful tool because it serves as a visual indicator of successful flow through the MVNs. We took advantage of this readout to characterize the effects of flow on MVNs with different starting geometries. We started with smaller or larger vessels, applied flow using identical pump settings, and evaluated MVNs expressing KLF2-GFP after 48 hours. While flow increased average vascular diameter of MVNs with both smaller and larger initial vessels, the diameters of MVNs under flow were indistinguishable, regardless of their starting point. Since the geometry of self-assembled engineered networks like ours depends on multiple conditions, (including cell counts, thrombin concentrations, fibrinogen efficacy, and time of gel injection into the device), which are prone to human and reagent variability, flow can serve as a means of reducing variability in samples across batches and users. While permeability tended to decrease with flow in both smaller and larger MVNs, it was striking that the lowest permeability was observed in the larger vessels group exposed to flow. While we evaluated permeability after 48 hours of flow, we did not characterize changes at earlier timepoints and it is possible that a period of increased permeability would be observed when the cells are undergoing junctional rearrangement^52^. In this case, MVNs that started out with smaller vessels would have to undergo more drastic structural changes to reach the 48-hour endpoint we observed and would reach their minimum permeability later than MVNs that started out with larger vessels and require far less structural adaptation.

The system that we have presented here, which incorporates MVNs with a KLF2-GFP reporter and a microfluidic pump, can be modified and expanded for future studies aimed at understanding the effects of flow on endothelial cell biology. For instance, MVNs could be formed from human pluripotent stem cell-derived ECs^7^ to study how shear stresses impact their phenotype and arteriovenous specification^53,54^. Further, alternative flow profiles could be explored using the KLF2-GFP readout, since it has been found that while KLF2 expression in response to high shear stress levels can be further increased by applying pulsatile flows^16^.

## Methods

### Cell culture and evaluation of monolayers

HUVECs were isolated and cultured^55^ and subsequently transduced with a pTRH-mCMV-dscGFP lentiviral reporter construct and sorted using fluorescence-activated cell sorting (FACS) in order to obtain cells with low GFP expression at baseline that will activate GFP in response to flow, as previously described^21^. KLF2-GFP HUVECs were maintained in VascuLife VEGF medium (Lifeline Cell Technology) prepared using the supplements as directed by the manufacturer, with the exception of heparin sulfate, which was added at a lower concentration (0.19 U/mL final concentration). KLF2-GFP HUVECs were grown to 90% confluency prior to use in MVNs and used at passage 14. Human lung fibroblasts (Lonza) at passage 7 were maintained in FibroLife S2 Fibroblast Medium (Lifeline Cell Technology) and cultured until 50-70% confluency prior to use in MVNs. All cells were maintained at 37°C and 5% CO_2_ throughout culture.

HUVEC monolayers were exposed to flow using a cone and plate device programmed to generate laminar flow and a specified shear stress (0.5 Pa and 1 Pa, equivalent to 5 dyn/cm^2^ and 10 dyn/cm^2^, respectively, in this study) and assayed for KLF2-GFP expression. As previously described^21^, KLF2-GFP HUVECs were plated on plastic substrates (QC Precision Machining) coated with 0.1% gelatin (Becton Dickinson) at a density of 6x10^4^ cells/cm^2^ 24 hours prior to the start of flow. One hour prior to flow initiation, the medium was replaced with new medium supplemented with 1.7% dextran (Catalog No. D5376, Millipore Sigma). Monolayers were maintained under flow for 24 hours at 37°C, 5% CO_2_ in a custom dynamic flow system^40^. Immediately after the cessation of flow, monolayers were imaged for GFP expression using an Axiovert microscope (Zeiss), and subsequently dissociated, stained with for VE-cadherin, and analyzed using FACS, as described below.

### Fabrication of microfluidic device and pump

Device and pump designs were created in Fusion 360 (Autodesk) and component molds were milled using a Computer Numerical Control (CNC) Milling Machine (Bantam Tools). Polydimethylsiloxane (PDMS, Dow Corning Sylgard 184, Ellsworth Adhesives) was mixed at a ratio of 10:1, degassed for 30 minutes, poured into milled molds, degassed a second time for 30 minutes, and cured in an oven at 65°F overnight. Microfluidic devices were subsequently fabricated as previously described^24,56^: individual PDMS devices were cut, gel and media ports were punched out using biopsy punches (Integra Miltex), and devices were sterilized by autoclave for 25 minutes. Devices and clean glass coverslips (VWR) were plasma treated (Expanded plasma cleaner, Harrick Plasma), bonded, and placed in an oven at 75°F overnight. Pump PDMS layers and membranes were trimmed, ports were punched out, and individual pumps were assembled after plasma bonding, as previously described^22^.

### Formation of microvascular networks

KLF2-GFP HUVECs and fibroblasts were suspended in a fibrin gel and seeded as previously described^6^. HUVECs and fibroblasts were dissociated from cell culture flasks using TrypLE Express (Thermo Fisher Scientific), suspended in a solution of VascuLife and 6 mg/mL fibrinogen from bovine plasma (Millipore Sigma) at a concentration of 14 million ml^-1^ and 2 million ml^-1^ HUVECs and fibroblasts, respectively. For each device, 7 μL of the cells and fibrinogen solution was combined with 7 μL of VascuLife with 2.8 U/mL thrombin from bovine plasma (Millipore Sigma) prior to injection in the device gel port (for a final concentration of 7 million ml^-1^ HUVECs, 1 million ml^-1^ fibroblasts, 1.4 U/mL of thrombin, and 3 mg/mL of fibrinogen). For some experiments, the thrombin concentration was decreased in order to slow gelling time, resulting in vessels with larger diameters, as indicated in the text. The remaining HUVECs were replated for seeding monolayers. Devices were incubated at 37°C for 10 minutes prior to adding VascuLife medium in the media channels. Device media was changed daily.

HUVEC monolayers were seeded along the sides of the gel channels of each device on day 4 of culture^6^. After removing the media, 30μg/mL fibronectin (Millipore Sigma) was introduced into each media channel and the devices were incubated for 30 minutes prior to HUVEC seeding. In each device, the fibronectin was removed from one media channel and a suspension of 0.5-1 million ml^-1^ HUVECs was added to the channel. The device was tilted so that the cell suspension would settle on top of the central gel channel and left for 10 minutes before repeating the procedure in the other media channel. Each device was then incubated for 10 minutes at 37°C prior to replacing the media in both channels.

### Application of flow in MVNs and measurement of flow speeds

The hydraulic resistance of the MVNs was measured by attaching 1mL syringe barrels to each media port of the devices, adding media to establish an initial pressure head difference of approximately 2.5 cm across the MVNs, and monitoring the changes in the pressure head difference at 15 minute intervals over 75 minutes. The media height difference between the two sides of the device over time could be fitted using the formula:

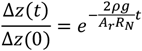

where Δz is the difference in media height, t is time, *ρ* is the density of cell culture media (∼993 kg/m^3^), *g* is the standard gravity (9.8 m/s^2^) and *R*_*N*_ is the bulk hydraulic resistance of the MVNs. *R*_*N*_ could then be calculated from:

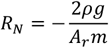

where m is the slope of the line fitted to the natural logarithm of the height difference.

The hydraulic MVN resistance was used in a combined lumped-element and finite-element model^22^ to predict the vascular wall shear stresses resulting from a range of pressures applied to the pump chamber. For flow experiments, pump pressures were selected based on these modeling results in order to achieve physiological vascular wall shear stress (0.5 Pa to 1.5 Pa). Each microfluidic device was attached to a pump and media was circulated through the microfluidic devices by applying cyclical air pressure through a solenoid and into the pumping chamber, as detailed previously^22^. Unless otherwise specified, flow was applied to MVNs for 48 hours prior to MVN evaluation.

To determine the flow speeds within vessels, MVNs were stained with 1:200 DyLight 594 UEA I lectin (Vector Laboratories) in VascuLife medium for 20 minutes at 37°C before 1μm fluorescent beads (Millipore Sigma) were added to the cell culture media in the microfluidic devices and pumps. Vessels and beads were visualized on an Axiovert microscope (Zeiss) and 60 fps video recordings of bead flows in response to application of 3-9 kPa of pressure at the pumping chambers were acquired. Pumps were run for 5 minutes at each pressure to reach stabile flow speeds before acquisition of 60 second videos. Custom MATLAB (MathWorks) scripts were used analyze bead speeds. First, videos were first divided into 60 frame (for 3kPa and 6kPa flows) or 30 frame (for 9kPa flows) segments. For each of these segments, maximum signal intensity projections over time were generated and the first frame was subtracted, leaving the bead tracks. These images were then thresholded to segment out the bead tracks, the connected components were determined, and the time at each point along each connected component was computed. The tracks were skeletonized and their length was calculated. Finally, the speed of each track was computed from the length and the start and end times of each component. Average bead speeds were determined by computing the mean of all of the bead tracks across all samples for each pump pressure.

### Imaging of MVNs and measurement of vascular permeability

Permeability of MVNs was measured by introducing fluorescent dextran (Texas Red, 70,000 MW, Thermo Fisher Scientific) into the media channels of each microfluidic device and quantifying the change in intravascular and extravascular fluorescence levels after 12 minutes, as previously described^6^. MVNs were imaged using an Olympus FV1000 confocal microscope at 37°C and using a 10x objective. Confocal z-stack images were acquired at a resolution of 0.5 pixels/μm at a z-spacing of 5 μm every 12 minutes. Image segmentation and average fluorescent intensity measurements were conducted using ImageJ and vascular permeability calculations were subsequently performed as detailed previously^6^.

Confocal z-stack images of KLF2-GFP signal were acquired as described for permeability above. Fiji was used to create maximum intensity projection images, apply Otsu thresholding to segment out GFP-positive signal, and measure the area and signal intensities of GFP-positive regions. GFP-positive areas are represented as a percentage of the total area of each image and GFP intensities are normalized to the lowest average intensity in a static sample of the same batch of MVNs.

To perform time course KLF2-GFP imaging, MVNs were stained with UEA I lectin as detailed above prior to the start of flow. Samples were maintained at 37°C, 5% CO_2_ in a custom enclosure while confocal z-stack images were acquired every 20 minutes as detailed above. GFP-positive area was determined using Fiji as detailed above, normalized from 0 to 1 for each time course, and fitted using an exponential curve to determine the time constant.

### Calculation of flow rates and wall shear stresses

Confocal images of static MVNs perfused with dextran were acquired as described above and tiled to span the entire width of the vasculature between the media channels. We used the micro-Vasculature Evaluation System algorithm to perform 3D vessel segmentation, skeletonization, and calculation of flow rates, speeds, and wall shear stresses in each branch^25^. For each MVN, boundary conditions were set such that the pressure across the network resulted in an average vessel flow speed of 2 mm/s. Wall shear stresses less than 0.01 Pa or greater than 5 Pa were omitted from data plots in order to remove vessels that did not experience flow or had negligibly small diameters.

### Analysis of gene expression

Cells were extracted from MVNs by manually cutting out the gel from the microfluidic devices and digesting each gel in 500 μL of 0.5 mg/mL Liberase TM (Millipore Sigma) in DMEM (Thermo Fisher Scientific) for 30 minutes on ice with intermittent agitation^56^. Cells from 6 devices were pooled, filtered to remove debris, stained with Alexa Fluor 647 Mouse Anti-Human CD144 antibody (Catalog No. 561567, BD Pharminogen) and sorted using fluorescence activated cell sorting (FACS) to isolate ECs for subsequent gene expression analysis. RNA was isolated by lysing cells using buffer RLT (Qiagen) and purified using RNeasy Micro Kit (Qiagen). Reverse transcription was performed using High-Capacity RNA-to-cDNA Kit (Thermo Fisher Scientific). Quantitative real-time polymerase chain reaction (qRT-PCR) was performed using Taqman PCR Master Mix (Applied Biosystems) and samples were amplified on an ABI Prism 7900HT Fast Detection System (Applied Biosystems). Gene expression was normalized against GAPDH endogenous controls.

### Analysis of vessel geometry

Total vascular area was calculated from maximum intensity projection images of MVNs perfused with dextran. Images were processed using AutoTube software^57^, with the following settings: pre-processing by adaptive histogram equalization, illumination correction, and image denoising through Block-Matching and 3D Filtering, tube detection with the Multi-Otsu thresholding, and tube analysis with removal of short ramification that are less than 20 pixels of length and merging of branch points within a radius of 22 pixels. Number of branches, branch length, and diameter were calculated by analyzing confocal z-stacks of MVNs perfused with dextran with the micro-Vasculature Evaluation System algorithm, which performs vessel segmentation, skeletonization, branch point analysis, and calculation of vascular morphological perameters in 3D^25^. To visualize vascular diameters, MVN surfaces were constructed from dextran images using Imaris software (Oxford Instruments) and clipped along the xz plane to reveal vessel openings.

### Perfusion of platelets in MVNs

Human whole blood was collected in sodium citrate and centrifuged for 15 minutes at 120g to separate the plasma and buffy coat layers from the red blood cells. Suspended platelets were labelled by incubating in 0.5μg/mL CD41 antibody (Catalog No. 303725, Biolegend) for 20 minutes at room temperature. Afterwards, the platelet solution was diluted 1:4 in PBS and recalcified by addition of 10mM CaCl_2_ immediately prior to perfusion through MVNs. To add platelets to the MVNs, media from one channel was removed, the channel was filled with platelet solution, media from the other channel was removed, and a platelet solution of equal volume to the solution in the other channel was added. For MVNs cultured under flow, the pumps were stopped and removed prior to addition of the platelets in the system. MVNs and platelets were imaged on a confocal microscope, just as described for permeability measurements. The percentage of CD41^+^ area in each image was determined from maximum intensity projections, using the same method described for determining the percentage area expressing KLF2-GFP described above.

### Statistics

Average values are reported as mean±standard deviation and samples sizes are reported in figure legends. Statistical analysis was performed using GraphPad Prism software. Parametric, two-tailed t tests were applied to analyses involving two groups and ordinary one way ANOVA with Tukey’s multiple comparison tests were applied to analyses involving more than two groups. * indicates p<0.05, ** indicates p<0.01, *** indicates p<0.001, **** indicates p<0.0001, and ns indicates p>0.05.

