## Supplementary Figures 1 and 2 for "Engineering microvascular networks using a KLF2 reporter to probe flow-dependent endothelial cell function"

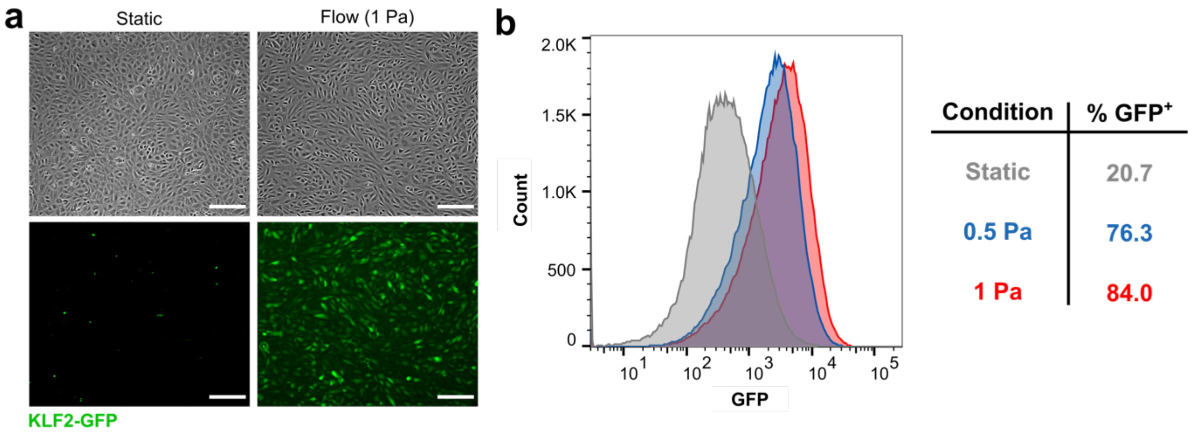


**Supplementary Figure 1. KLF2-GFP expression in HUVEC monolayers under flow. a** Example of HUVEC monolayer in static culture and under flow for 24 hours. Monolayers were imaged using brightfield microscopy (top row) and GFP fluorescence to detect expression of KLF2 (bottom row). Scale bars = 200 μm. **b** Flow cytometry of static monolayers, and monolayers cultured under flow at 0.5 Pa or 1 Pa of shear stress.

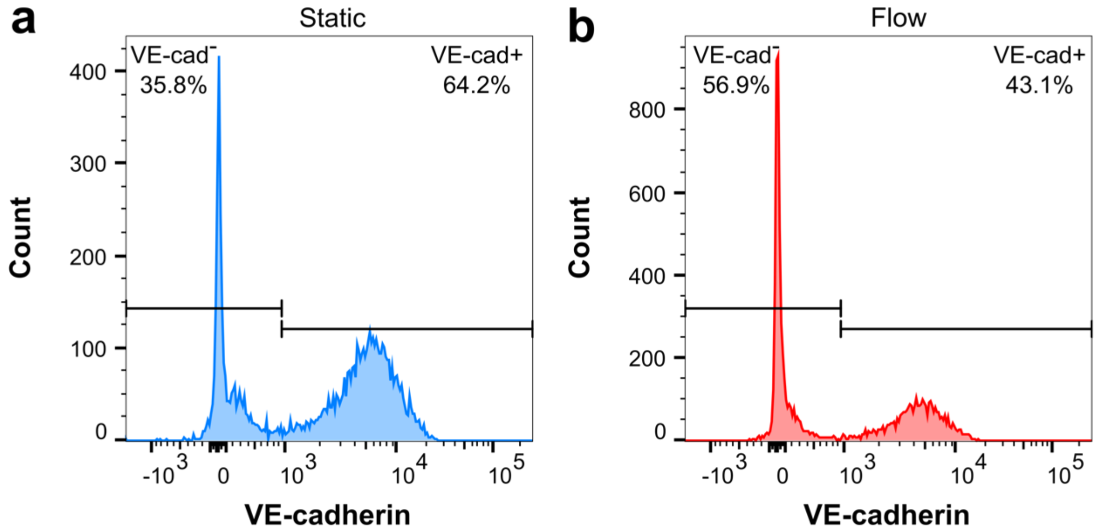


**Supplementary Figure 2. Separation of fibroblasts and endothelial cells in MVNs based on VE-cadherin expression. a** Flow cytometry count of cells in static MVNs. **b** Flow cytometry count of cells in flow MVNs.
